## Supplementary figures for "The non-catalytic role of DNA polymerase epsilon in replication initiation in human cells"

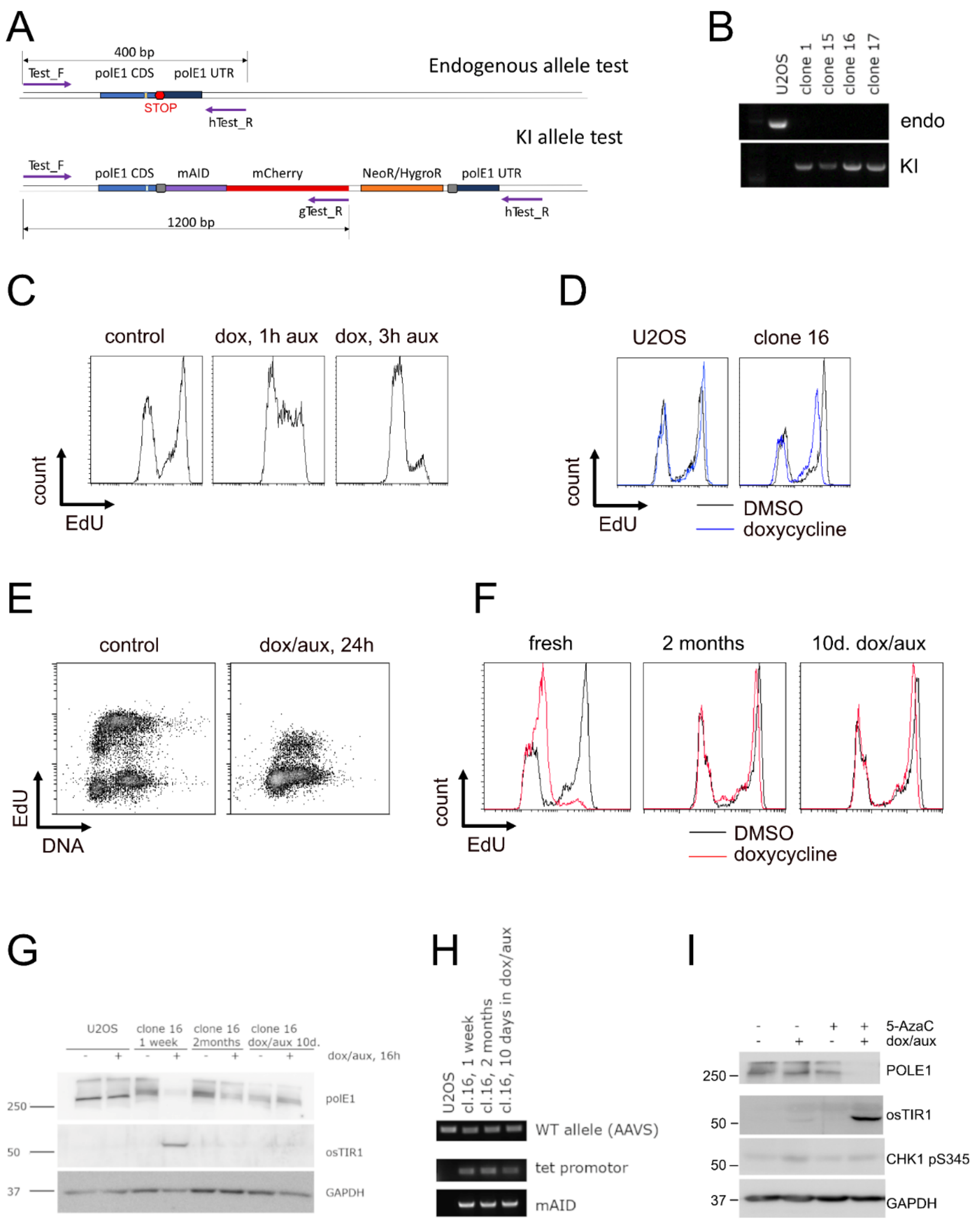


**Figure S1. A.** Schematic representation of the PCR based testing of the knock-in efficiency. **B.** PCR testing the knock-in clones. Genomic DNA from control U2OS cells or indicated clonal lines was used as a template for PCR with primers Test_F and hTest_R (for endogenous allele) or Test_F and gTest_R (for knock-in allele). **C.** Clone 16 cells were treated for 16h with doxycycline, followed by 1h or 3h treatment with auxin, as indicated. EdU was added for the last 30 min. Flow cytometry analysis of EdU incorporation is shown. **D.** U2OS, clone 16 were treated for 24h with doxycycline. EdU was added for the last 30 min. Flow cytometry analysis of EdU incorporation is shown. **E.** Homozygous mAID-KI clone 15 cells were treated for 24h with DMSO or dox/aux, 10µM EdU was added for the last 30 min of treatment**.** Flow cytometry plots showing EdU incorporation and DNA content (7-AAD staining) are shown. **F-H.** Clone 16 cells: fresh, cultured for 2 months without dox/aux, or cultured for 2 weeks with dox/aux were treated with DMSO or dox/aux for 16h. **F.** 10µM EdU was added for the last 30 min of treatment**.** Flow cytometry plots showing EdU incorporation are shown. **G.** Western blots of the total cell lysates are shown. **H.** DNA gel of the PCR of genomic DNA with primers against the indicated regions is shown. **I.** Clone 16 cells cultured for 2 months were treated with 5-AzaC for 48h, dox/aux or DMSO were added for the last 16h of treatment where indicated. Western blots of the total cell lysates are shown.

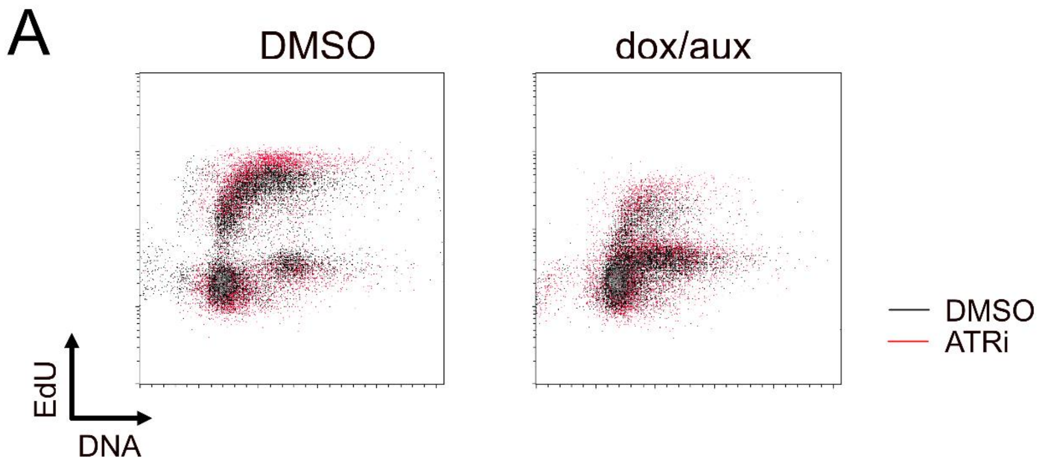


**Figure S2.** Clone 16 cells were treated for 16h with DMSO or dox/aux as indicated, DMSO or 5µM ATRi was added to the indicated samples for 60 min before harvest, 10µM EdU was added for the last 30 min of treatment. Flow cytometry plots showing EdU incorporation and DNA content are shown.

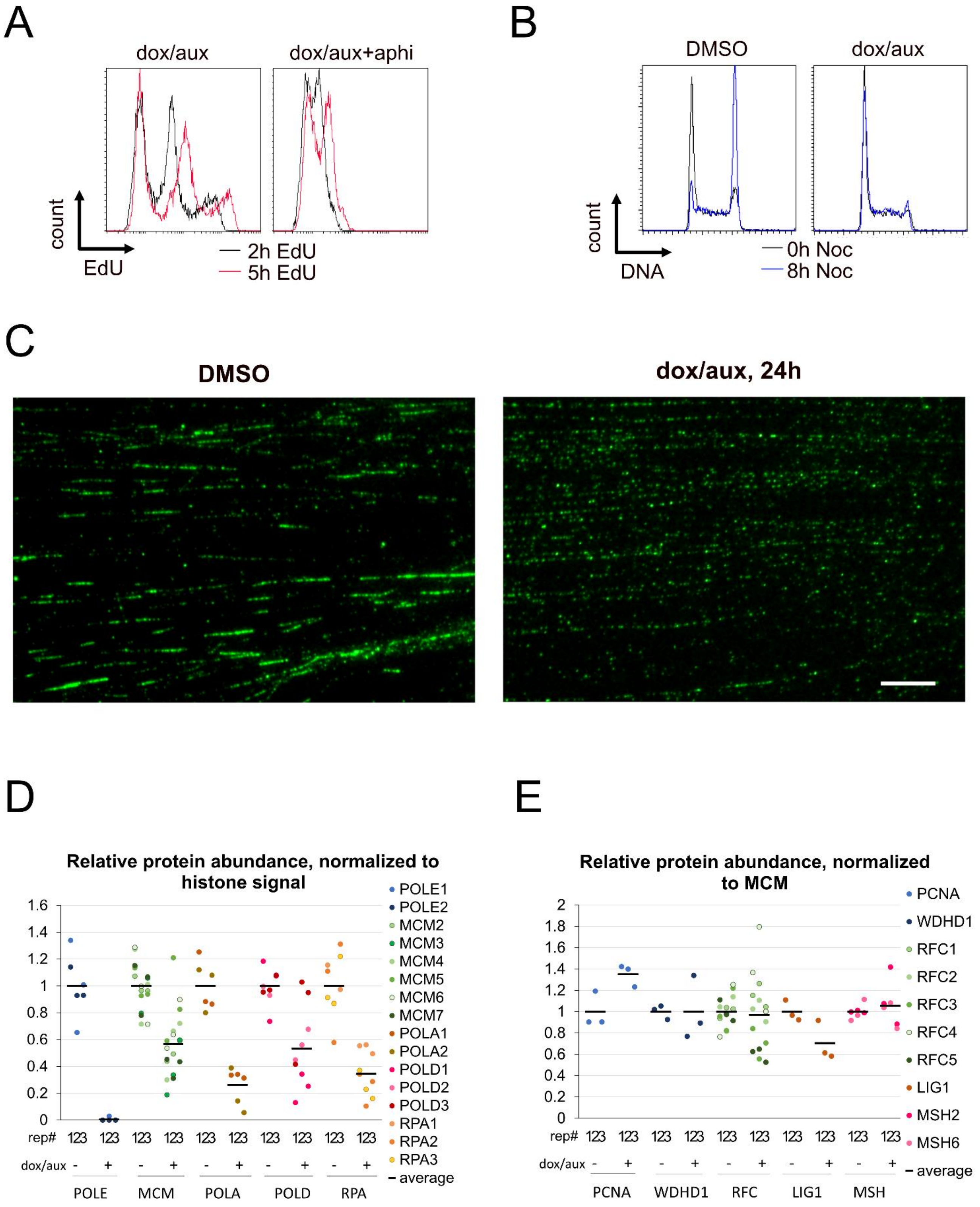


**Figure S3. A.** Clone 16 cells were treated for 16h with dox/aux, 10µM EdU was added for the indicated times before harvest, followed by ethanol fixation. 2µM aphidicolin was added 1h before the start of the EdU pulses where indicated. Flow cytometry plots showing EdU incorporation are shown. **B.** Clone 16 cells were treated for 16h with DMSO or dox/aux, followed by 8h nocodazole treatment where indicated. Flow cytometry plots of DNA content (PI) are shown. **C.** Clone 16 cells were treated for 24h with DMSO or dox/aux. Ongoing replication was labeled with 10 min pulse of IdU and visualized using DNA fiber analysis, as described in Methods. Scale bar is 10 µm. **D-E.** Clone 16 cells were treated for 16h with DMSO or dox/aux, followed by 10 min EdU pulse and iPOND isolation of protein, associated with nascent DNA, and mass-spectrometry. The signal was normalized to average signal of histones (D) or MCM subunits (E) in each sample, and to respective DMSO-treated samples. The data from three experimental repeats are shown.

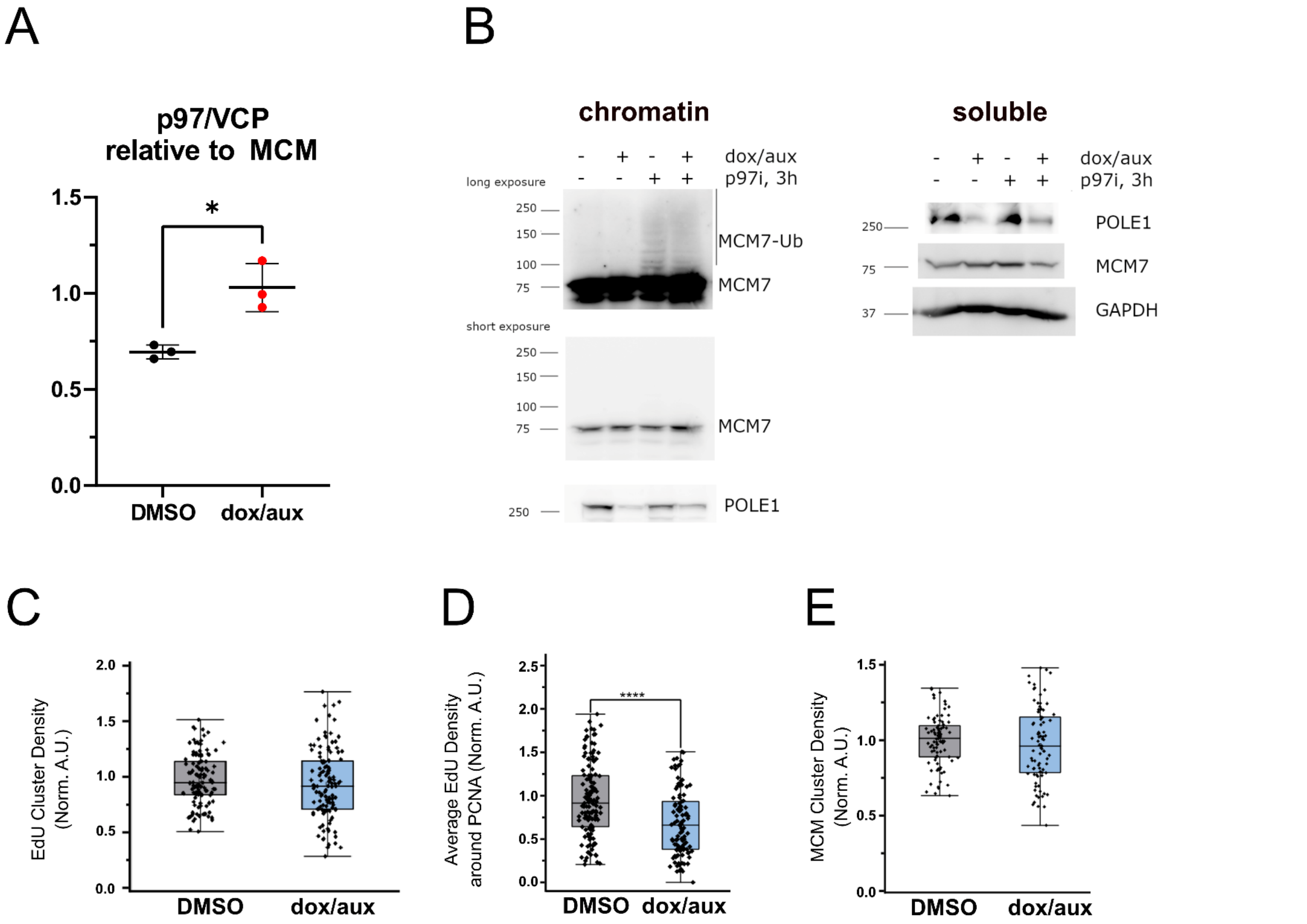


**Figure S4. A.** Clone 16 cells were treated for 16h with DMSO or dox/aux, followed by 10 min EdU pulse and iPOND isolation of protein, associated with nascent DNA, and mass-spectrometry. The signal was normalized to average signal of MCM subunits in each sample. The data from three experimental repeats are shown. P-value was calculated using paired t-test, based on three experimental repeats. **B.** Clone 16 cells were treated for 16h with DMSO or dox/aux, followed by 3h p97i treatment where indicated. Western blots of soluble land nuclease insoluble chromatin fractions are shown. **C-E.** Clone 16 cells treated for 16 h with DMSO or dox/aux were pulse labeled with thymidine analogue for 15 minutes prior to processing for super-resolution imaging. Quantitation of number of EdU clusters (C), average density of EdU around PCNA (D) and number of MCM clusters (E) detected within a 6 × 6 μm$^2$ square region of interest, normalized to DMSO treated clone 16 cells based on at least 2 independent experiments. (For EdU cluster density, n= 118, 118; average EdU density around PCNA, n= 136, 109; MCM cluster density, n= 94, 86 for DMSO and dox/aux treated clone 16 cells, **** P < 0.0001, student's t-test).

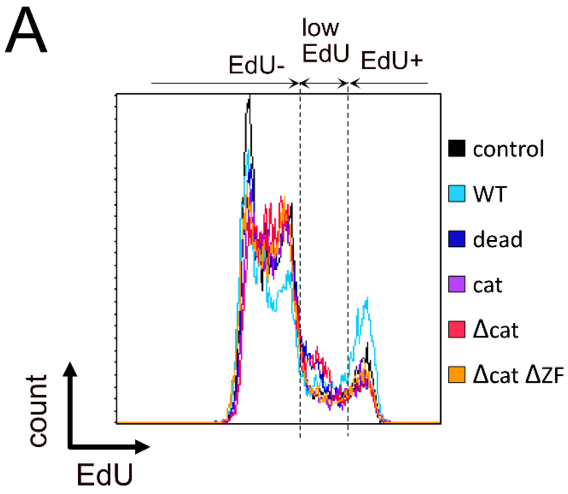


**Figure 5S. A.** Clone 16 cells were transfected with indicated constructs. 32h later dox/aux were added to the cells for 16h. 10µM EdU was added for the last 30 min. Flow cytometry histograms of the EdU channel and gating for “EdU-“, “low EdU” and “high EdU” fractions are shown.

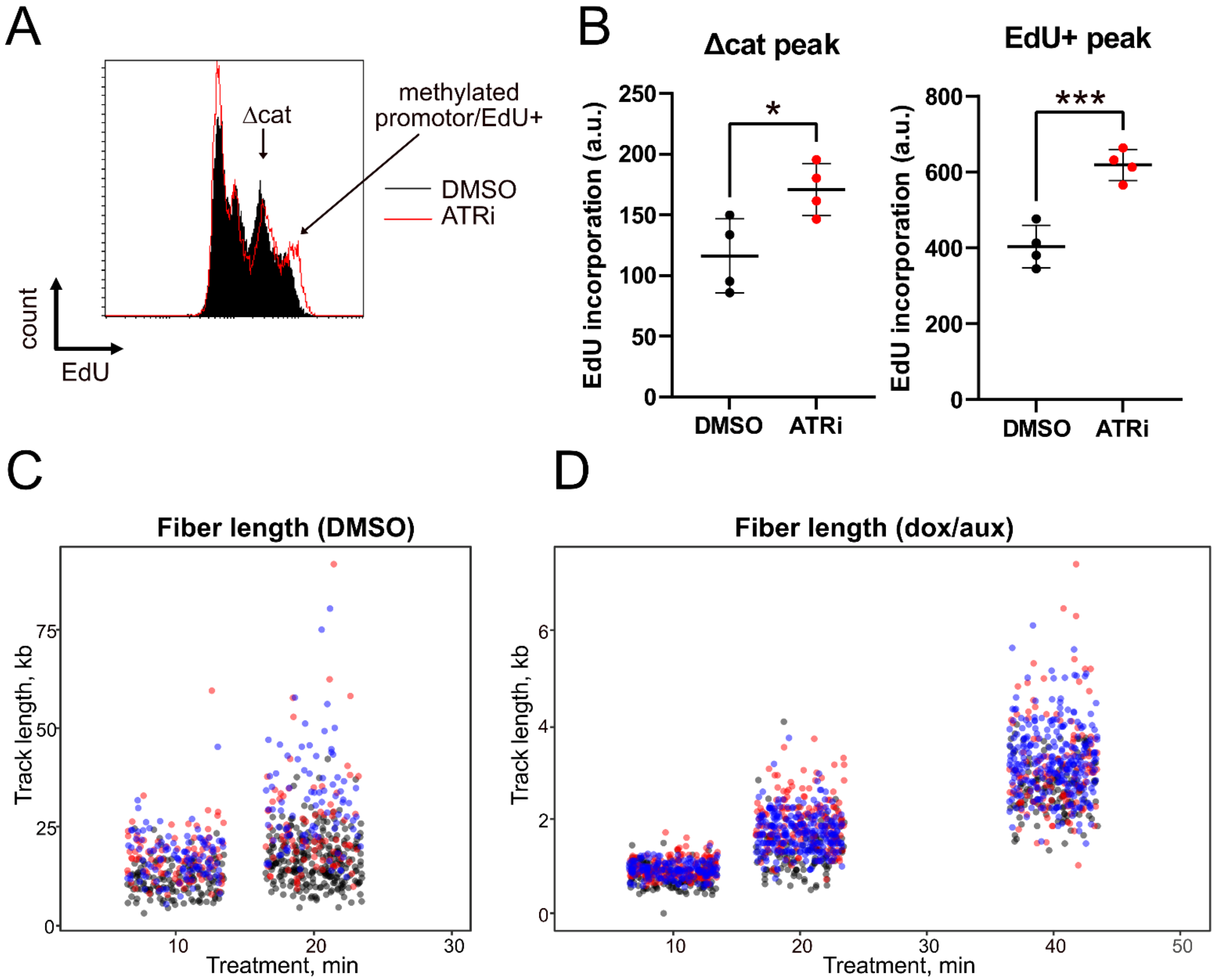


**Figure S6. A-B.** Cells stably expressing myc-FLAG-Δcat (1.6+Δcat) were treated for 16h dox/aux, DMSO or 5 μM ATRi were added 15 min before the start for the 30 min EdU pulse. Flow cytometry histograms of EdU incorporation (A) or EdU incorporation quantifications (B) are shown. Quantification is based on three independent experiments, means and standard deviations are shown, dox-resistant population was counted as EdU+, t-test was used for statistical analysis. **P < 0.001, *P < 0.05. **C-D.** 1.6+Δcat cells were treated for 1 6h with DMSO (C) or dox/aux (D). Ongoing replication was labeled with 10- or 40-min pulse of CldU followed by 20 min pulse of IdU and visualized using DNA fiber analysis (same experiments as Fig. 6E), as described in Methods. Individual measurements of fiber lengths are shown (three colors represent three experimental repeats).
